# *Distribution of GOPC:ROS1* and other *ROS1* fusions in glioma types

**DOI:** 10.1101/2021.04.30.442173

**Authors:** Philipp Sievers, Damian Stichel, Martin Sill, Daniel Schrimpf, Dominik Sturm, Florian Selt, Jonas Ecker, Evelina Miele, Mariëtte E. G. Kranendonk, Bastiaan B. J. Tops, Patricia Kohlhof-Meinecke, Rudi Beschorner, Christof M. Kramm, Martin Hasselblatt, Guido Reifenberger, David Capper, Pieter Wesseling, Till Milde, Andrey Korshunov, Olaf Witt, Stefan M. Pfister, Wolfgang Wick, Andreas von Deimling, David T. W. Jones, Felix Sahm

**Author notes:** Corresponding author: Felix Sahm, MD, PhD Department of Neuropathology University Hospital Heidelberg and Clinical Cooperation Unit Neuropathology (B300) German Cancer Research Center (DKFZ) Im Neuenheimer Feld 224 69120 Heidelberg, Germany Fon. Co-first author (Philipp Sievers, Damian Stichel). **Authorship statement** Experimental design: P.S., D. Stichel, S.M.P, A.v.D., D.T.W.J., F. Sahm Implementation: P.S., D. Stichel, D.T.W.J., F. Sahm Data analysis/interpretation: P.S., D. Stichel, M.S., D.T.W.J., F. Sahm Sample collection: P.S., D. Sturm, F. Selt, J.E., E. M., M.E.G.K., B.B.J.T., P. K-M., R. B., C. M. K., M. H., G. R., D. C., P.W., T.M., A. K., O. W., S. M. P., W. W., A.v.D., D.T.W.J., F. Sahm Approved final manuscript: All authors.

## Abstract

The ROS proto-oncogene 1 (*ROS1*) gene is rearranged in various cancers. The translated fusion protein presents an attractive therapeutic target, since specific inhibitors have been approved for several tumor types. In glioma, *ROS1* fusions are frequent within infantile hemispheric glioma, and single case reports on occurrences in other glioma types exist. However, a comprehensive analysis spanning the full width of glioma types and subtypes is lacking. We here assessed the spectrum and distribution of *ROS1* fusions by screening >20,000 glioma cases for typical chromosomal alterations, with subsequent RNA-sequencing for confirmation of candidate cases. *ROS1* fusions were identified in 16 cases, from low grade pilocytic astrocytoma WHO grade 1 to glioblastoma, IDH wildtype WHO grade 4. Thus, despite being enriched in some tumor types, *ROS1* fusions are not pathognomonic for specific glioma types and may consitute a relevant target in a variety of cases.

## Introduction

Gliomas are the most common primary tumors of the central nervous system (CNS). Among low-grade gliomas, mitogen-activated protein kinase (MAPK) pathway alterations are frequent and may provide a therapeutic target. Currently, mechanism-of-action based therapeutic approaches outside the MAPK pathway are scarce. However, especially patients with subtotally resected, recurrent or highly malignant tumors may substantially benefit from the identification of additional specific oncogenic drivers that not only provide insight into disease pathogenesis but also offer targets for personalized cancer therapies. The ROS proto-oncogene 1 (*ROS1*) gene encodes a receptor tyrosine kinase that is involved in chromosomal rearrangements in various cancers^1^, which present an attractive therapeutic target, since specific inhibitors have been approved for several entities^2,3^. Data on *ROS1* fusions in glioma are limited to single cases or small series^4–7^.

Recently, an enrichment of these fusions was found in a small number of mostly gliomas in infants^8,9^. Routine diagnostic assessment of *ROS1* status in gliomas, however, is so far restricted to a few specialized centers or molecularly informed trials^10^. Thus, the landscape of *ROS1* fusions across a broad series of glial tumors of all age groups has not been comprehensively studied so far. Consequently, the distribution among the various types of low- to high-grade glioma is unknown. Similarly, no data exists to determine whether *ROS1* fusion-positive gliomas, irrespective of histology, may share further biological features, potentially supporting a ‘ROS1-subtype’ of gliomas. Here, we investigated the presence of *ROS1* fusions in a large cohort of 20,723 patients encompassing different diagnostic entities within the spectrum of glioma, to elucidate the frequency of such fusions and the characteristics of the respective cases.

## Methods and Results

To identify gliomas with structural alterations affecting chromosome 6q (around the *ROS1* locus), we systematically evaluated copy-number data of our DNA methylation dataset encompassing 20,723 gliomas, irrespective of specific entity and WHO grade (Suppl. Fig. 1 and 2). As a high proportion of *ROS1* fusions (in particular the most frequent *GOPC:ROS1* fusion) are accompanied by a segmental loss of chromosome 6q22 in the copy-number profile, DNA methylation data were screened for a segmental loss covering that region (Suppl. Fig. 1). Automated analysis was followed by visual inspection and led to the identification of 14 potential cases. On suspicious cases, we performed RNA and targeted exome sequencing, and confirmed the presence of *ROS1* fusions in all 14 tumors (Fig. 1A). In the most common (n=11) *GOPC:ROS1* fusions (Fig. 1B), exons 1-7 or 1-4 of *GOPC* (NM_001017408) are fused in frame to exons 35-43 of *ROS1* (NM_002944). Single cases of exons 36-43 of ROS1 fused downstream of *ZCCHC8* exons 1-2 (NM_0017612), *ARCN1* exons 1-5 (NM_001655), or *CHCHD3* exons 1-2 (NM_017812) were also observed (Fig. 1C). In all fusion events, the kinase domain of *ROS1* was retained (Fig.1B). In addition, two further *ROS1*-fused glioma samples that were already detected as such by performing RNA sequencing in a diagnostic context, after the initial screen was performed were included into subsequent analyses. One of the samples harbored a *GOPC:ROS1* fusion (with exons 1-7 of *GOPC* fused to exons 35-43 of *ROS1*) and indeed showed segmental loss of chromosome 6q22, while the other case harbored a *CEP85L:ROS1* fusion (with exons 1-12 of *CEP85L* (NM_001042475) fused exons 35-43 of *ROS1*) with a segmental gain of chromosome 6q22. In addition, we analyzed RNA sequencing data from a set of > 1000 FFPE tissue samples processed in a diagnostic setting. Here, no further gliomas harboring a *ROS1*-fusion were detected.

**Fig. 1:**
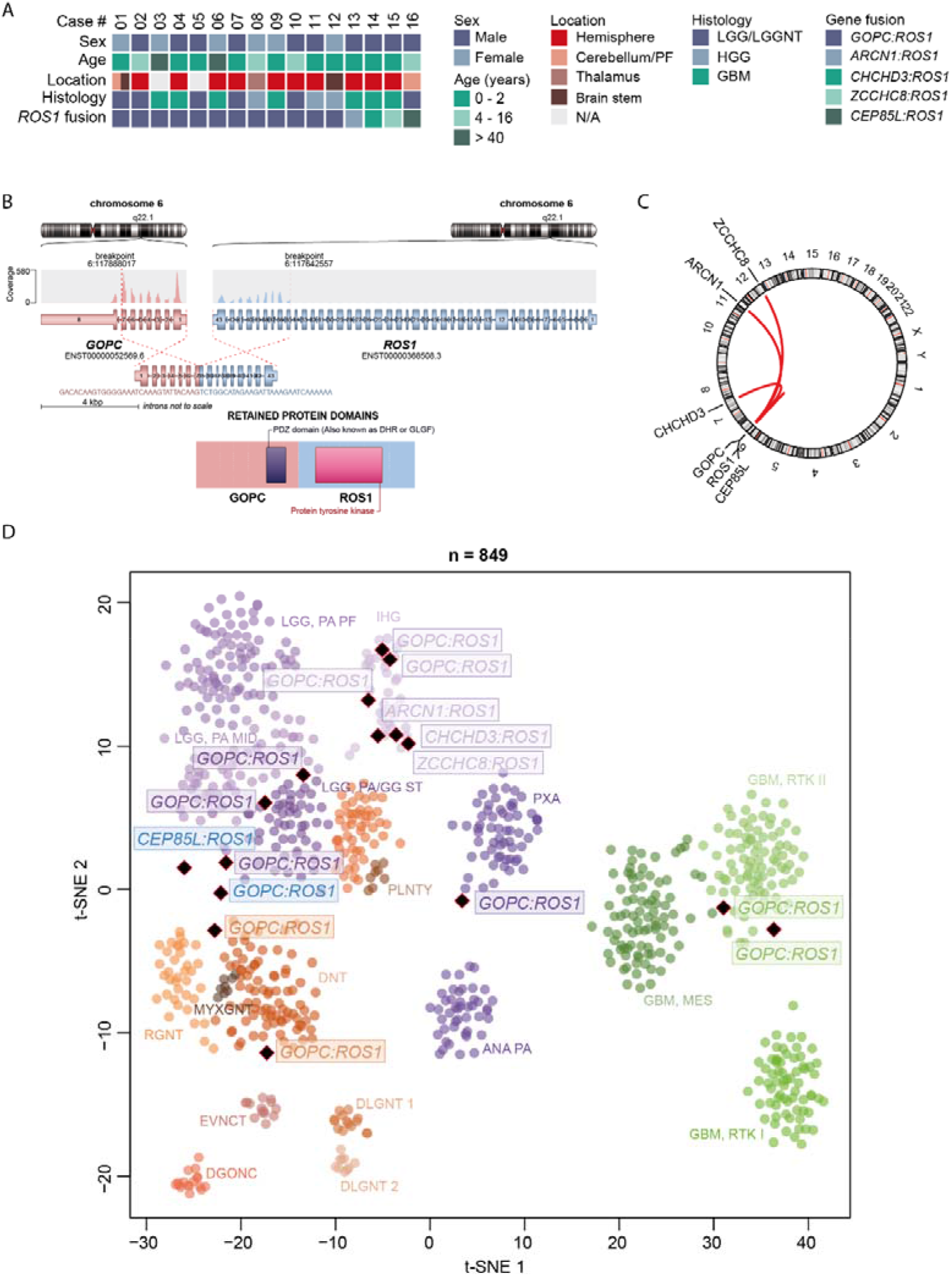
Summary of clinico-pathological characteristics and key molecular findings in tumors with *ROS1* gene fusion (A). Schematic illustration of the *GOPC:ROS1* fusion detected in case #3 involving exons 1-7 of *GOPC* and exons 35-43 of *ROS1* (B). Circos plot of gene fusions targeting *ROS1* (Lines link fusion gene partners according to chromosomal location; C). t-distributed stochastic neighbor embedding (t-SNE) analysis of DNA methylation profiles of *ROS1*-fused glioma alongside selected reference samples (D). Reference DNA methylation classes: posterior fossa pilocytic astrocytoma (LGG, PA PF), hemispheric pilocytic astrocytoma and ganglioglioma (LGG, PA/GG ST), midline pilocytic astrocytoma (LGG, PA MID), polymorphous low-grade neuroepithelial tumor of the young (PLNTY), ganglioglioma (LGG, GG), diffuse leptomeningeal glioneuronal tumor subgroup 1 (DLGNT 1), diffuse leptomeningeal glioneuronal tumor subgroup 2 (DLGNT 2), infantile hemispheric glioma (IHG), extraventricular neurocytoma (EVNYCT), dysembryoplastic neuroepithelial tumor (LGG, DNT), rosette-forming glioneuronal tumor (LGG, RGNT), myxoid glioneuronal tumor of the septum pellucidum and lateral ventricle (MYXGNT), diffuse glioneuronal tumor with oligodendroglioma-like features and nuclear clusters (DGONC), anaplastic astrocytoma with piloid features (ANA PA), pleomorphic xanthoastrocytoma (PXA), glioblastoma IDH wildtype subclass RTK I (GBM, RTK I), glioblastoma IDH wildtype subclass RTK II (GBM, RTK II), glioblastoma IDH wildtype subclass mesenchymal (GBM, MES). The two *ROS1*-fused glioma samples that were already detected as such by performing RNA sequencing in a diagnostic context are highlighted in blue. Other abbreviations: LGG/LGGNT, low-grade glioma/low-grade glioneuronal tumor; HGG, high-grade glioma; GBM, glioblastoma; PF, posterior fossa; N/A, not available.

A t-distributed stochastic neighbor embedding (t-SNE) analysis of DNA methylation profiles alongside a broad reference set of CNS tumors^11^ revealed that the ‘ROS1 cohort’ molecularly segregated into different glioma groups (Fig. 1D). Six of the samples grouped with the DNA methylation class infantile hemispheric glioma, other tumors clustered with various reference classes of glioma from low- to high-grade (Fig. 1D). Histological re-evaluation confirmed the different histological entities and underline that *ROS1* fusions are not specific to any one glioma entity. Interestingly, most of the patients harboring a fusion were children (particularly infants). Of note, however, was the finding that two classical adult IDH-wildtype glioblastomas in adult patients also harbored a *GOPC:ROS1* fusion.

## Discussion

Our data show a high frequency of *ROS1* gene fusions within the DNA methylation class infantile hemispheric glioma, which is in line with recent studies^8,9^. This clinically distinct group of gliomas (that were initially often diagnosed as glioblastomas) carries a high prevalence of gene fusions with *ROS1*, *ALK*, *NTRK1/2/3*, or *MET* as a fusion partner. However, our finding that *ROS1* fusions also occur in cases that were both histologically and epigenetically clearly pilocytic astrocytoma or IDH-wildtype glioblastoma, respectively, underscores that this event is not pathognomonic for infantile hemispheric glioma, nor limited to pediatric patients, so in that respect concerns a quite ‘promiscuous’ marker in that respect.

Although relatively rare in other gliomas, identification of *ROS1* fusions is important from a treatment perspective, as there are specific inhibitors available. Screening via copy-number profiling and subsequent validation using RNA sequencing provides an efficient approach to identify patients who may benefit from this targeted therapy. However, as illustrated by one of the cases that was identified by performing RNA sequencing in a diagnostic setting, not all variants of *ROS1* fusion necessarily show a deletion around the *ROS1* locus. For example, copy-neutral translocations can lead to *ROS1* fusions as well, and such cases would be missed by screening for segmental 6q22 loss. RNA sequencing thus remains the ‘gold standard’ for adequate detection of these rare events.

Our findings highlight *ROS1* fusions as a rare but potentially highly relevant therapeutic target for a subset of patients with gliomas of different histological grades and biological classes. Even though these fusions have no strong diagnostic relevance, since they are not pathognomonic for a tumor type, they are in line with the increasing demand to provide predictive markers in diagnostic neuropathology. This highlights the need for expanded testing for such alterations beyond infant gliomas. It will be interesting to see whether ROS1-inhibitors will be effective in upcoming clinical trials for glioma patients.

## Acknowledgements

We thank J. Meyer, L. Hofmann and A. Habel for skillful technical assistance and the microarray unit of the DKFZ Genomics and Proteomics Core Facility for providing Illumina DNA methylation array-related services.

