## Supplementary material for "*Distribution of GOPC:ROS1* and other *ROS1* fusions in glioma types": Suppl. Data

**Supplementary materials and methods**

Sample collection

Tumor samples and patient data were obtained from multiple national and international collaborating centers and collected at the Department of Neuropathology of the University Hospital Heidelberg (Germany). Analysis of tissue and clinical data was performed in accordance with local ethics regulations. Clinical details of the patients are listed in Supplementary Table 1.

DNA methylation array processing and copy number profiling

Genomic DNA was extracted from fresh-frozen or formalin-fixed and paraffin-embedded (FFPE) tissue samples. Genome-wide DNA methylation profiling of all samples was performed using the Infinium MethylationEPIC (EPIC) BeadChip (Illumina, San Diego, CA, USA) or Infinium HumanMethylation450 (450k) BeadChip array (Illumina) according to the manufacturer’s instructions and as previously described^1^. Raw data were generated at the Department of Neuropathology of the University Hospital Heidelberg, the Genomics and Proteomics Core Facility of the German Cancer Research Center (DKFZ) or at respective international collaborator institutes, using both fresh-frozen and formalin-fixed paraffin-embedded (FFPE) tissue samples. All computational analyses were performed in R version 3.6.0 (R Development Core Team, 2016; <https://www.R-project.org>). Copy-number variation analysis from 450k and EPIC methylation array data was performed using the conumee Bioconductor package version 1.12.0^2^. Raw signal intensities were obtained from IDAT-files using the minfi Bioconductor package version 1.21.4. Illumina EPIC and 450k samples were merged to a combined data set by selecting the intersection of probes present on both arrays (combineArrays function, minfi). Each sample was individually normalized by performing a background correction (shifting of the 5% percentile of negative control probe intensities to 0) and a dye-bias correction (scaling of the mean of normalization control probe intensities to 10,000) for both color channels. Subsequently, a correction for the array type (450k/EPIC) was performed by fitting univariable, linear models to the log2-transformed intensity values (removeBatchEffect function, limma package version 3.30.11). The methylated and unmethylated signals were corrected individually. Beta-values were calculated from the retransformed intensities using an offset of 100 (as recommended by Illumina). All samples were checked for duplicates by pairwise correlation of the genotyping probes on the 450k/EPIC array. To perform unsupervised non-linear dimension reduction, the remaining probes after standard filtering^1^ were used to calculate the 1-variance weighted Pearson correlation between samples. The resulting distance matrix was used as input for t-SNE analysis (t-distributed stochastic neighbor embedding; Rtsne package version 0.13). The following non-default parameters were applied: theta = 0, pca = F, max_iter = 10,000 perplexity = 20.

Automated screening and identification of *ROS1*-fused glioma samples using DNA methylation data

For identification of putative relevant cases showing a *ROS1* fusion we pre-analyzed DNA methylation data of 20,723 glioma samples as follows. As *ROS1* fusions are typically accompanied by a segmental loss of chromosome 6q22 in the copy-number profile, DNA methylation data from 20,723 glioma samples were screened for a segmental loss of this region. Cases with low quality copy-number profiles were filtered. Automated analysis was followed by visual inspection and led to identification of 12 cases with sufficient material for confirmation via RNA sequencing.

RNA sequencing and analysis

RNA was extracted from FFPE tissue samples using the automated Maxwell system with the Maxwell 16 LEV RNA FFPE Kit (Promega, Madison, WI, USA), according to the manufacturer’s instructions. Transcriptome analysis using messenger RNA (mRNA) sequencing of samples was performed on a NextSeq 500 instrument (Illumina) as previously described^3^. Fastq files from transcriptome sequencing were used for *de novo* annotation of fusion transcripts using the deFuse^4^ and Arriba (v1.2.0)^5^ algorithms with standard parameters. The fusion plot was created using the R-script for visualization provided in Arriba.

DNA sequencing and mutational analysis

Capture-based next-generation DNA sequencing was performed on a NextSeq 500 instrument (Illumina) as previously described^6^ using a custom brain tumor panel (Agilent Technologies, Santa Clara, CA, USA) covering the entire coding and selected intronic and promoter regions of 130 genes of particular relevance in central nervous system tumors.

References to supplementary methods

**Supplementary Table 1 - Summary of clinicopathological characteristics and key genetic alterations**

| **Case** | **Sex** | **Age (years)** | **Location** | **Histology** | **MC prediction v12** | **Gene fusion** | **Indication for a *ROS1* fusion by CNV** | **Additional CNV** | **NGS findings** |
| --- | --- | --- | --- | --- | --- | --- | --- | --- | --- |
| **#01** | **f** | **1** | **cerebellum, brain stem** | **LGG** | **LGG, PA PF** | ***GOPC:ROS1*** | **focal loss chr 6q** | **-** | **-** |
| **#02** | **m** | **6** | **parietal left** | **LGG/LGGNT** | **LGG, PA/GG** | ***GOPC:ROS1*** | **focal loss chr 6q** | **unclear baseline** | **-** |
| **#03** | **f** | **45** | **n/a** | **GBM** | **GBM, RTK II** | ***GOPC:ROS1*** | **focal loss chr 6q** | **gain chr 7, loss chr 10, gain chr 20, loss chr 22q, CDKN2A/B homozygous deletion, CDK4 amplification, MDM4 amplification** | **TERT promoter C250T**  **PTEN:NM_000314:exon2:c.G141T:p.R47S** |
| **#04** | **m** | **0** | **hemispheric left** | **GBM** | **IHG** | ***GOPC:ROS1*** | **focal loss chr 6q** | **CDKN2A/B homozygous deletion** | **-** |
| **#05** | **m** | **2** | **n/a** | **pO** | **nc** | ***GOPC:ROS1*** | **focal loss chr 6q** | **seg. loss chr 22q** | **-** |
| **#06** | **f** | **68** | **fronto-parietal left** | **GBM** | **GBM, RTK II** | ***GOPC:ROS1*** | **focal loss chr 6q** | **gain chr 7, loss chr 10, CDKN2A/B homozygous deletion** | **TERT promoter C250T**  **TP53:NM_001126116:exon1:c.G71C:p.R24P** |
| **#07** | **m** | **2** | **left ventricle** | **PMA** | **nc** | ***GOPC:ROS1*** | **focal loss chr 6q** | **seg. loss chr 19p** | **-** |
| **#08** | **f** | **12** | **thalamic right** | **GNT (high-grade)** | **nc** | ***GOPC:ROS1*** | **focal loss chr 6q** | **loss chr 1p, CDKN2A/B homozygous deletion** | **-** |
| **#09** | **f** | **0** | **hemispheric left** | **GBM** | **IHG** | ***GOPC:ROS1*** | **focal loss chr 6q** | **-** | **-** |
| **#10** | **m** | **7** | **right ventricle** | **PA** | **nc** | ***GOPC:ROS1*** | **focal loss chr 6q** | **gain chr 14q, loss chr 22q** | **-** |
| **#11** | **m** | **0** | **hemispheric right** | **AA** | **IHG** | ***GOPC:ROS1*** | **focal loss chr 6q** | **-** | **n/a** |
| **#12** | **f** | **2** | **medulla oblongata** | **HGG** | **nc** | ***GOPC:ROS1*** | **focal loss chr 6q** | **-** | **-** |
| **#13** | **m** | **2** | **fronto-parietal right** | **GBM** | **IHG** | ***ARCN1:ROS1*** | **focal loss chr 6q** | **loss chr 10, seg. loss chr 2q, 5p, 9p, 11q** | **PTEN:NM_000314:exon5:c.C388G:p.R130G** |
| **#14** | **m** | **0** | **fronto-parietal right** | **GBM** | **IHG** | ***CHCHD3:ROS1*** | **focal loss chr 6q** | **gain chr 11q, loss chr 6p** | **-** |
| **#15** | **m** | **2** | **temporo-parietal right** | **GBM** | **IHG** | ***ZCCHC8:ROS1*** | **focal loss chr 6q** | **gain chr 8** | **-** |
| **#16** | **m** | **16** | **posterior fossa** | **PA** | **nc** | ***CEP85L:ROS1*** | **focal gain chr 6q** | **-** | **-** |

Abbreviations: f, female; m, male; n/a, not available; LGG, low-grade glioma; LGGNT, low-grade glioneuronal tumor; GBM, glioblastoma; pO, pediatric oligodendroglioma; PMA, pilomyxoid astrocytoma; GNT, glioneuronal tumor (high-grade); AA, anaplastic astrocytoma; PA, pilocytic astrocytoma; MC, DNA methylation class; LGG, PA PF, posterior fossa pilocytic astrocytoma; LGG, PA/GG, hemispheric pilocytic astrocytoma and ganglioglioma; GBM, RTK II, glioblastoma IDH wildtype, subclass RTK II; IHG, infantile hemispheric glioma; nc, not classifiable; CNV, copy number variations; NGS, next-generation sequencing.

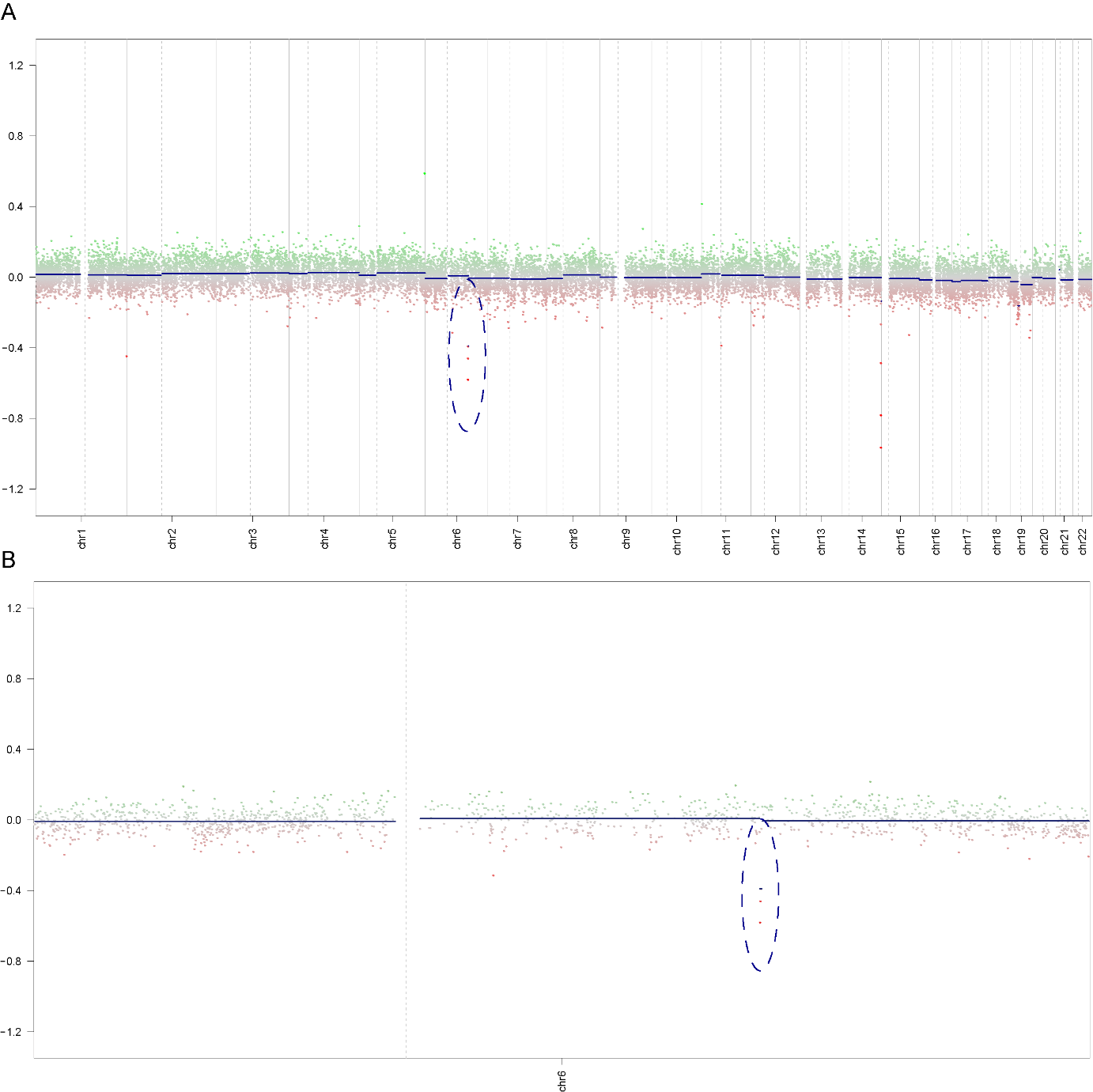

**Supplementary Fig. 1** Copy-number profile derived from DNA methylation array data showing a focal loss of chromosome 6q22 around the *ROS1* locus (A). Enlarged view of chromosome 6 from the same case (B).

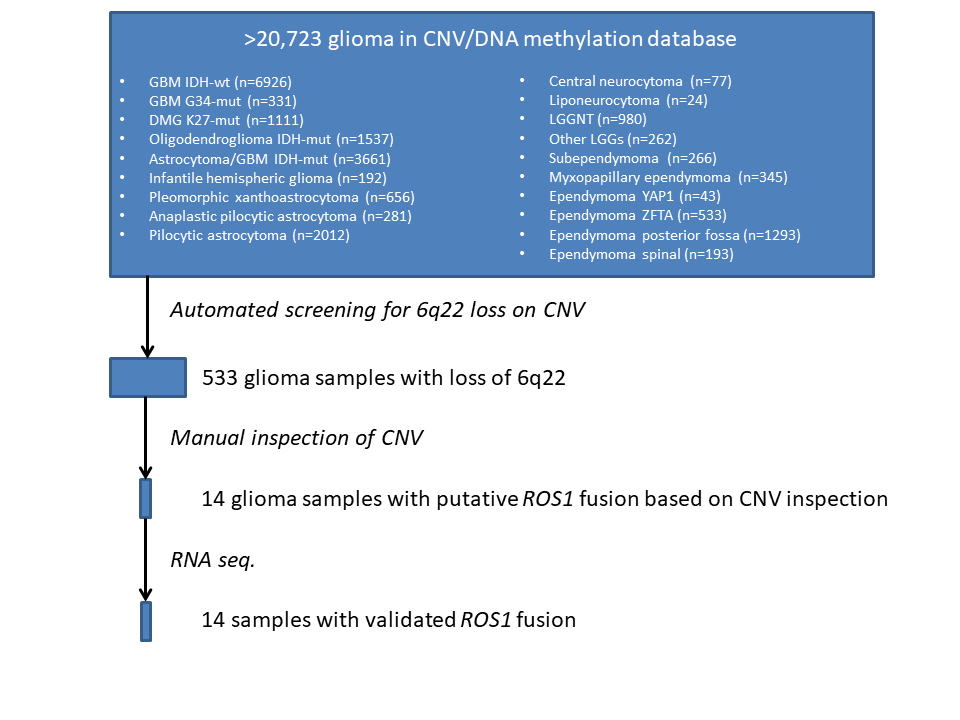
**Supplementary Fig. 2** Workflow for identification of the *ROS1*-fused gliomas.

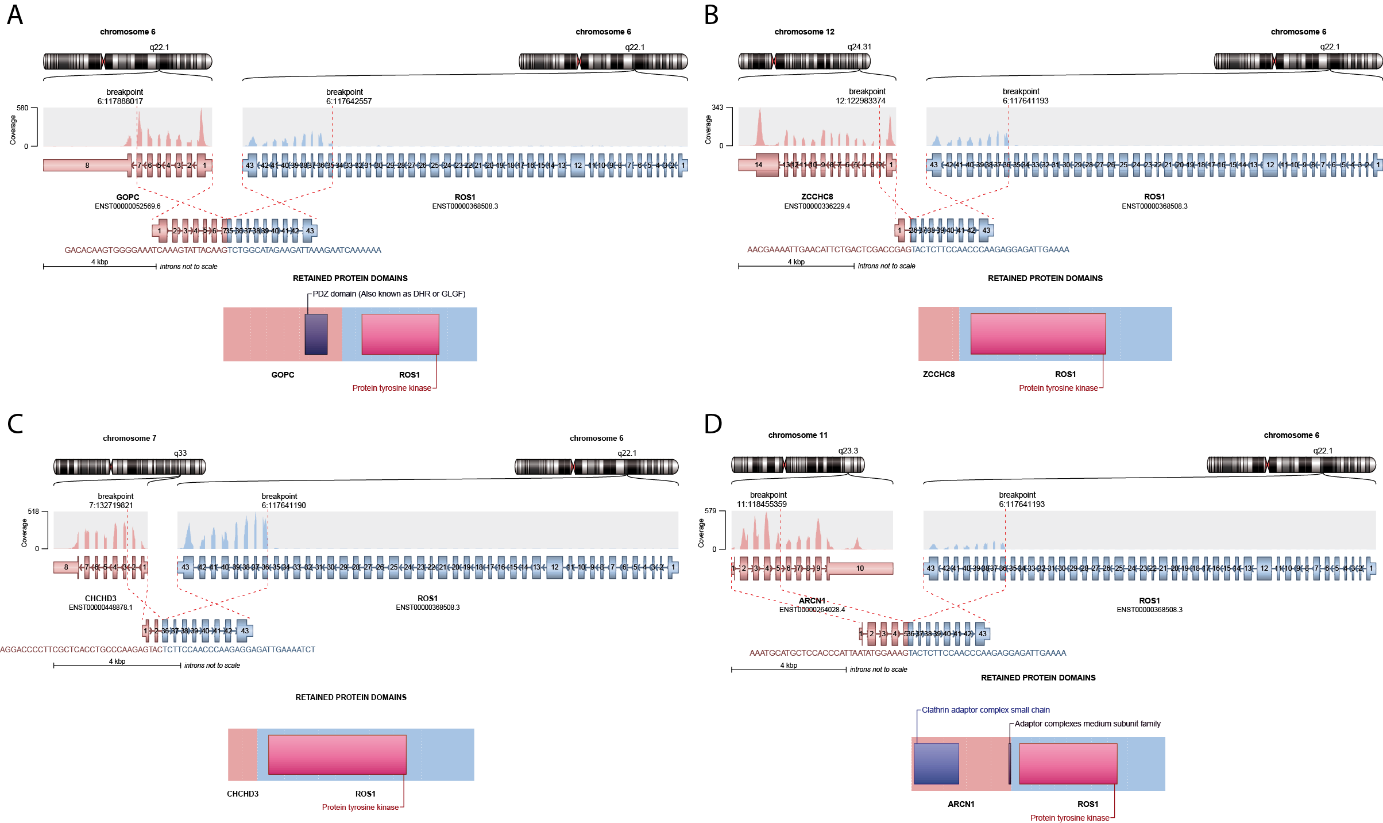

**Supplementary Fig. 3** Schematic illustration of the different *ROS1*-fusions detected in the glioma series - *GOPC:ROS1-* (A), *ZCCHC8:ROS1*- (B), *CHCHD3:ROS1*- (C) and *ARCN1:ROS1*-fusion (D).
